# TMPRSS2 reduces antibody recognition of SARS-CoV-2 spike

**DOI:** 10.64898/2026.02.17.706397

**Authors:** Andrea Cottignies-Calamarte, Alejandro De Cruz, Cyril Planchais, Françoise Porrot, Mateo Kryzpow, Martin Jungbauer-Groznica, Eva Thuillier, Amelie Willeveau, Isabelle Staropoli, Florence Guivel-Benhassine, Pierre Rosenbaum, Ignacio Fernández, Felix Rey, Julian Buschrieser, Sophie Trouillet-Assant, Hugo Mouquet, Olivier Schwartz, Timothée Bruel

## Abstract

The serine protease TMPRSS2 acts as a cofactor for SARS-CoV-2 entry by cleaving the viral spike (S) to initiate fusion. Whether TMPRSS2 has an impact on humoral immune response against S remains poorly characterized. Here, we show that TMPRSS2 impairs antibody binding to S. In S-expressing and infected cells, TMPRSS2 decreases monoclonal antibody (mAb) and immune serum binding, as well as antibody-dependent cellular cytotoxicity (ADCC) induction. Using a panel of 39 mAbs targeting various S regions, we observe that those binding to the S2 subunit are the most affected by TMPRSS2. TMPRSS2 promotes a partial shedding of S1 and changes S2 conformation. This processing reduces Angiotensin-Converting Enzyme 2 (ACE2) binding while increasing cell-cell fusion. We further observe that the capacity of TMPRSS2 to decrease antibody recognition is conserved across coronaviruses and shared with other TMPRSS proteins. However, TMPRSS2 expression in infected cells does not impact significantly virions’ infectivity or the antibody recognition, as measured by flow virometry. Collectively, our findings suggest that TMPRSS2 processing of S favors a fusion intermediate conformation which is less sensitive to antibody recognition.

## Introduction

The SARS-CoV-2 Spike (S) is a trimeric glycoprotein, responsible for viral fusion. Each protomer is composed of two subunits, S1 and S2, separated by a furin cleavage site (FCS). The S1 subunit comprises the receptor binding domain (RBD), which interacts with the cellular receptor Angiotensin-Converting Enzyme 2 (ACE2), and the N-terminal domain (NTD) (Hoffmann et al., 2020). S2 encompasses the fusion machinery and the transmembrane domain (Hills et al., 2025; Lusvarghi et al., 2025; Marcink et al., 2022; Zhang et al., 2025). Interaction between the RBD and ACE2 leads to S1 shedding and unmasking of the S2 fusion peptide (FP) (Grunst et al., 2024; Lusvarghi et al., 2025; Marcink et al., 2022). Insertion of the FP within the target membrane requires a proteolytic cleavage in a second site within S2, named S2’, which is remarkably conserved across coronaviruses. Processing of the S2’ site occurs at the plasma membrane or in endosomes, by transmembrane serine protease 2 (TMPRSS2) or cathepsins, respectively (Kakizaki et al., 2024). TMPRSS2 may also process S at the FCS site, as furin proteases and TMPRSS2 cleavage signals overlap (R–X–[K/R]–R↓ and [K/R]↓-S, respectively) (Fraser et al., 2022). Other members of the TMPRSS family can process S protein at S1/S2 and S2’ sites, albeit with variable efficacy (Kishimoto et al., 2021; Zang et al., 2020). Proteolytic cleavage by TMPRSS2, and possibly other proteases, may occur outside of the canonical S1/S2 and S2’ sites with functional consequences that remain largely misunderstood (Fraser et al., 2022; Lusvarghi et al., 2025; Meng et al., 2022). Thus, cellular proteases shape SARS-CoV-2 S protein structure and function.

Despite intracellular assembly and budding of viral particles, SARS-CoV-2-infected cells express S at the plasma membrane due to leaky coatomer interaction (Dey et al., 2022). If neighboring cells express ACE2 it may lead to syncytia formation, where TMPRSS2 appears to be dispensable though it accelerates the process (Buchrieser et al., 2020; Cattin-Ortolá et al., 2021). Membrane-bound S expression also enables the clearance of infected cells via antibody Fc-mediated functions, such as ADCC (Dufloo et al., 2021; Guenthoer et al., 2024), which may participate in the protection mediated by non-neutralizing anti-S antibodies (Clark et al., 2024). Yet, the main correlate of protection against COVID-19 is antibody neutralization (Earle et al., 2021; Goldblatt et al., 2022; Khoury et al., 2021). The most potent neutralizing antibodies target the RBD, although antibody responses are also elicited against other S regions (Bean et al., 2025; Clark et al., 2024; Guenthoer et al., 2024; Ju et al., 2020; Pinto et al., 2020; Planchais et al., 2022; Zhou et al., 2023; Zost et al., 2020). Both vaccination and infection elicit neutralizing antibodies, but the emergence of immune-evasive variants enables viral circulation in immune populations (Dufloo et al., 2021; Guenthoer et al., 2024; Natarajan et al., 2021; Ng et al., 2022, 2021; Planchais et al., 2022; Sterlin et al., 2021). Consequently, the SARS-CoV-2 pandemic is characterized by sequential waves of immune-evasive variants (Jian et al., 2024; Planas et al., 2024, 2022, 2021). Viral immune escape is mostly driven by mutations in key neutralizing epitopes. Other mechanisms, such as differential glycosylation and reduced S exposition by infected cells, also contribute to SARS-CoV-2 evasion of humoral immune responses (Boson et al., 2021; Kim et al., 2023; Sztain et al., 2021). Characterizing those alternative antibody evasion strategies requires an in-depth understanding of S-antibody interactions. The S adopts various conformation (closed, opened, pre-fusion, post-fusion), which are not all equally recognized by antibodies. Structural approaches have extensively characterized extreme states (i.e. stabilized pre- and post-fusion trimers), but the landscape of intermediate conformations and their sensitivity to antibody is poorly characterized. The extend to which some of those conformations may contribute to antibody evasion is also unknown.

In this study, we hypothesize that the processing of SARS-CoV-2 S by TMPRSS2 will modify its recognition by antibodies. We observe that proteolytic cleavage of S by TMPRSS2 reduces antibody binding at the surface of infected cells. Not all antibodies are affected similarly, with those targeting S2 being the most sensitive. We further determine that TMPRSS2 rearranges S conformation to reduce antibody binding and ADCC induction against S-expressing cells while preserving S fusogenicity. In contrast, on viral particles, TMPRSS2 only moderately reduced antibody binding. Altogether, our results demonstrate that TMPRSS2 changes S conformation to impair antibody recognition and ADCC during SARS-CoV-2 infection, shedding light on a novel pro-viral role of this protease.

## Methods

### Antibodies and chemicals

Anti-SARS-CoV-2 spike monoclonal antibodies (mAbs) were produced in vitro as previously described (Planchais et al., 2022). Briefly, cDNA encoding for heavy and light antibody chains were either generated in-house from S-specific memory B cells (Planchais et al., 2024, 2022) or obtained from public sequences. All mAbs were produced as human IgG1 by co-transfection of 293-Freestyle (ThermoFisher), followed by affinity purification with protein G Sepharose 4 fast flow beads (GE Healthcare) and validated by ELISA for antigen specificity (Planchais et al., 2022). See **Table 1** for a detailed characterization of mAbs. TMPRSS2 staining was performed with an anti-TMPRSS2 nanobody described previously (Saunders et al., 2023). Hygromycin, puromycin and blasticidin were obtained from InvivoGen.

**Table 1:** List of anti-spike antibodies used in this study. n.d.: Not determined; N/A: Not applicable

|  |  |  |  |  |
| --- | --- | --- | --- | --- |
| Spike region | International Nonproprietary Names | Code mAb | Epitope | Reference |
| Anti-RBD | Bamlanivimab | BAM | Class Ia | Jones B. et al., Sci Transl Med, 2021 |
|  | Imdevimab | IMD | Class Ia | Hansen J. et al., Science, 2020 |
|  | Cilgavimab | CIL | Class Ia | Zost S.J. et al., Nature, 2020 |
|  | Sotrovimab | S309 | Class Ia | Pinto D. et al., Nature, 2020 |
|  | Tixagevimab | TIX | Class Ia/Ib | Zost S.J. et al., Nature, 2020 |
|  | Bebtelovimab | LyCoV-1404 | Class Ia/Ib | Westendorf K. et al., Cell Rep, 2022 |
|  | Casirivimab | CAS | Class Ib | Hansen J. et al., Science, 2020 |
|  | N/A | SA55 | Class I | Cao Y. et al., Cel Rep, 2022 |
|  | Etesevimab | ETE | Class Ia/IIa | Shi R. Nature, 2020 |
|  | Adintrevimab | ADG2 | Class I/IV | Rappazzo C.G. et al., Science, 2021 |
|  | Regdanvimab | REG | Class IV | Kim C. et al., Nat Comm, 2021 |
|  | N/A | 2-43 | Trimer interface | Liu L. et al., Nature, 2020 |
| Anti-NTD | N/A | Cv2.5105 | Non-supersite | Planchais C. et al. *J Exp Med* 2022 |
|  | N/A | Cv2.1153 | Non-supersite | Planchais C. et al. *J Exp Med* 2022 |
|  | N/A | Cv2.1191 | Non-supersite | Planchais C. et al. *J Exp Med* 2022 |
|  | N/A | Cv2.3144 | Non-supersite | Planchais C. et al. *J Exp Med* 2022 |
|  | N/A | Cv2.1309 | Non-supersite | Planchais C. et al. *J Exp Med* 2022 |
|  | N/A | Cv2.2333 | Non-supersite | Planchais C. et al. *J Exp Med* 2022 |
|  | N/A | Cv2.2396 | Non-supersite | Planchais C. et al. *J Exp Med* 2022 |
|  | N/A | Cv2.1177 | Supersite | Planchais C. et al. *J Exp Med* 2022 |
|  | N/A | Cv2.2221 | Supersite | Planchais C. et al. *J Exp Med* 2022 |
|  | N/A | Cv2.3215 | Supersite | Planchais C. et al. *J Exp Med* 2022 |
|  | N/A | Cv2.3296 | Supersite | Planchais C. et al. *J Exp Med* 2022 |
|  | N/A | Cv2.8145 | Supersite | Planchais C. et al. *J Exp Med* 2022 |
|  | N/A | Cv2.8162 | Supersite | Planchais C. et al. *J Exp Med* 2022 |
|  | N/A | Cv2.2157 | Supersite | Planchais C. et al. *J Exp Med* 2022 |
|  | N/A | Cv2.1177 | Supersite | Planchais C. et al. *J Exp Med* 2022 |
|  | N/A | Cv2.1169 | n.d. | Planchais C. et al. *J Exp Med* 2022 |
| Anti-S2 | N/A | Cv2.2180 | n.d. | Planchais C. et al. *J Exp Med* 2022 |
|  | N/A | Cv2.2216 | n.d. | Planchais C. et al. *J Exp Med* 2022 |
|  | N/A | Cv2.8187 | n.d. | Planchais C. et al. *J Exp Med* 2022 |
|  | N/A | Cv2.4102 | n.d. | Planchais C. et al. *J Exp Med* 2022 |
|  | N/A | Cv2.1380 | n.d. | Planchais C. et al. *J Exp Med* 2022 |
|  | N/A | Cv2.3175 | n.d. | Planchais C. et al. *J Exp Med* 2022 |
|  | N/A | Cv2.4171 | n.d. | Planchais C. et al. *J Exp Med* 2022 |
|  | N/A | Cv2.9116 | n.d. | Planchais C. et al. *J Exp Med* 2022 |
|  | N/A | Cv2.3132 | Heptad repeat 2 (HR2) | Planchais C. et al. *iScience* 2024 |
|  | N/A | CoV44-62 | Fusion peptide | Dacon C. et al. *Science* 2022 |
|  | N/A | 76E1 | Fusion peptide | Sun X. et al. *Nat. Microbiol.* 2022 |
|  | N/A | C77G12 | Fusion peptide | Low S. et al. *Science* 2022 |
| n | 39 |  |  |  |

### Human sera

Serum of individuals with hybrid immunity (Wuhan infection followed by 3 mRNA vaccine doses; **Table 2**) were obtained in the framework of the Covid-Ser cohort. A written informed consent was obtained from all participants; ethics approval was obtained from the national review board for biomedical research in April 2020 (Comité de Protection des Personnes Sud Méditerranée I, Marseille, France; ID RCB 2020-A00932-37; NCT04341142).

**Table 2:** Patients characteristics of which serum were used for this study. Individuals received Pfizer-BioNTech COVID-19 vaccine (BNT162b2) and were admitted in Lyon Hospice Civils for COVID-19 infection. Autoimmune and inflammatory disease include: Adisson’s disease, Celiac disease, and Crohn’s disease.

|  |  |
| --- | --- |
| n | 15 |
| Age at blood sampling (years), median [IQR] | 55 [28–61] |
| Female, *n* (%) | 12 (80) |
| Time elapsed between the last vaccination or infection event and blood sampling (days) | 110 [94–112] |
| Body mass index, *n* | 15/15 |
| Body mass index, median [IQR] | 23.88 [20.20–27.72] |
| Currently smoker, *n* (%) | 1/15 (6.66) |
| Alcohol consumption (daily), *n* (%) | 0/15 (0) |
| **Comorbidities** |  |
| Presence of comorbidities, *n* (%) | 3/15 (20) |
| Neurological disorders, *n* (%) | 1 (6.66) |
| Cardiovascular disorders, *n* (%) | 1 (6.66) |
| Hypertension, *n* (%) | 0 (0) |
| Heart failure, *n* (%) | 0 (0) |
| Diabetes, *n* (%) | 0 (0) |
| Immune deficiency, *n* (%) | 0 (0) |
| Liver disease, *n* (%) | 0 (0) |
| Kidney disease, *n* (%) | 0 (0) |
| History of cancer, *n* (%) | 0 (0) |
| Hypothyroidy, *n* (%) | 0 (0) |
| Rheumatic disease, *n* (%) | 1 (6.66) |
| Chronic lung disease, *n* (%) | 1 (6.66) |
| Autoimmune and inflammatory disease, *n* (%) | 0 (0) |

### Plasmids

SARS-CoV-2 spikes from D614G, Delta, BA.4/5, BQ1.1, XBB.1.5/9, EG.5.1, XEC, and KP3.1.1 were human codon-optimized and synthetized (GeneArt, Thermo Fisher Scientific) as described (Buchrieser et al., 2020). Downstream cloning was performed using the Gateway strategy (ThermoFisher) in phCMV and pLV-EF1a-IRES-Puro or Hygro (Addgene #85132 and #85134). Expression plasmid were all previously described: Myc- or mScarlet-I-tagged TMPRSS in pCAGS or pCDEST respectively (Hoffmann et al., 2021; Saunders et al., 2023), S from coronaviruses 229E, RatG13 and NL63 in pcDNA3 (Ng et al., 2022), pVAX expressing S-BANAL20-236 (Temmam et al., 2022), lentivector plasmid pLV-ACE2 (Buchrieser et al. 2020) and pQCXIP-GFP1-10 and pQCXIP-GFP11 (Addgene #68716 and #68715, respectively). All sequences were confirmed by sequencing (Eurofins Genomics). VSV-G pseudovirus vectors were produced by phosphate calcium transfection as previously described (Buchrieser et al., 2019).

### Cells

HEK293T, IGROV-1 and Caco2 cells were obtained from ATCC and maintained in DMEM glutamax (Gibco) containing 10% heat-inactivated FCS (Gibco) and 1% Penicilin-Streptomycin (Gibco). The absence of mycoplasma was monitored weekly using the MycoAlert kit (Lonza). Mucilair® primary human nasal epithelial cells (hNECs) were purchased from Epithelix (Switzerland). 293T-S and IGROV-1-TMPRSS2 were cultivated in presence of 1µg/ml puromycin or 250 µg/ml hygromycin, respectively, to maintain transgene expression. Caco2-S were generated by lentiviral transduction and selected with 1µ/ml puromycin 4-5 days before each experiment, because of the cytotoxicity of S in those ACE2-expressing cells. CD16 reporter Jurkat cell line was cultivated as per the manufacturer recommendations (Promega). The SARS-CoV-2 reporter cells S-Fuse (293T-GFP1-10 and 293T-GFP11) and IGROV-1-TMPRSS2 were previously described and maintained accordingly (Buchrieser et al., 2020, 2019; Planas et al., 2024).

### Viral production, titration and cell infection

Amplification and production of SARS-CoV-2 D614G and XBB.1.5 variants have been previously described (Planas, 2024). For S surface staining, 10^6^ Caco2, IGROV-1 (WT or TMPRSS2+) were plated in 6-wells plate. The next day, cells were infected with D614G or XBB1.5 variants for 2h in fresh media, after what media was changed. At 24h post-infection (hpi), Caco2 cells were either treated with Camostat (SigmaAldrich) or DMSO containing media and incubated for another 24h. IGROV-1 cells were infected for 48h. After infection, cells were washed 2 times in PBS EDTA and detached with trypsin before transferring to 96-wells plate for spike staining as described in the staining section. After fixation, cells were permeabilized for staining with Dylight488 anti-nucleocapsid (N) mouse IgG1 (in-house production). For D614G production in IGROV-1 WT or overexpressing TMPRSS2, 2x10^6^, cells were plated the day before infection in a T25 flask (Greiner). The next day, cells were infected and inoculum was replaced with fresh media after 2h. Supernatant was harvested at 2-and 3-days post-infection (dpi) and clarified at 1000g for 10min before storage at -80°C. Samples were incubated 10min at 80°C before quantifying N gene in the supernatants using the Luna universal RT-qPCR kit (New-England Biolab), the N2 primers tandem (Lu et al., 2020) and analyzed using a QS6 pro (ThermoFisher), as reference to a commercial standard (Sigma, EURM-001).

### Transfection

For TMPRSS2 transfection, 10^6^ cells in 3ml DMEM 10%FCS were plated in 6-wells plate and transfected with 1 µg plasmid DNA in 1ml Optimem (Gibco) and 12 µl Lipofectamin2000 (ThermoFisher). In multiple transfection conditions, 6.10^5^ cells were plated in 150 µl DMEM 10% FCS onto which 50µl Optimem containing 100ng DNA and 0.6 µl Lipofectamin2000 were added. Transfection lasted overnight before any downstream analysis or assay.

### Antibody binding analysis by flow cytometry

Cells were detached in PBS 1%EDTA or 0.05% trypsin (Gibco) and washed before staining with 1µg/ml final antibody concentration for 15min at 4°C in DMEM 10% FCS. Sera were used at a dilution of 1:100 in DMEM 10%FCS. Cells were then washed in PBS before staining with a goat anti-human IgG AlexaFluor 647 (Invitrogene) for 15min at 4°C, washed in PBS and stained with Fixable Aqua Dead cells (ThermoFisher) for 10min at 4°C. Next, cells were washed in DMEM 10%FCS and cells were fixed in PBS-4%PFA (ThermoFisher) for 15min before a final PBS wash. Flow cytometry analysis was performed on Attune machine (ThermoFisher) and analyzed on FlowJo V10.10.0 (BD Life Science). When indicated % reduction in binding was calculated based on the mean fluorescence intensity (MFI) as follows

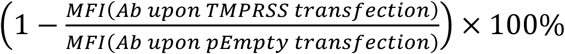

### S1 ELISA

Supernatants were centrifuged 5min 1.000g before storage at -20°C. Anti-NTD antibody Cv2.1169 was coated overnight in PBS at 5µg/ml onto 96 well plate (Corning #3361). Samples were diluted in PBS 5% FCS and incubated for 2h at 37°C after plating. Anti-RBD antibody SA55-biotinylated was diluted in PBS 5% FCS and incubated for 1h at 37°C, followed by streptavidin-HRP (BD, #554066, diluted 1000 times). Detection was performed using OPD (Sigma) and stopped using 3M sulfuric acid. Between each step, plates were thoroughly washed with PBS.

### Donor-Acceptor spike-mediated fusion

5x10^5^ 293T-S cells expressing GFP1-10 were cocultured in a 1:1 ratio with 293T-GFP11 stably expressing ACE2 or not in μClear black 96-well plate (Greiner Bio-One). Cells were incubated for 4h before fixation in 4% PFA final. Plates were then washed and nuclei were stained in PBS-Hoechst 33342 (Invitrogen; diluted 10.000 times) and images were acquired using the Opera Phenix High-Content Screening System (PerkinElmer). GFP area per well was calculated using Harmony 191 High-Content Imaging and Analysis Software (PerkinElmer, HH17000012, v.5.0). Normalized GFP area was calculated by dividing GFP area of the tested condition by GFP area observed in the absence of any TMPRSS2 transfection.

### ADCC assay

5x10^5^ transfected 293T-S-D614G were plated in white 96-wells plates with 100-fold diluted serum and 5x10^5^ CD16 reporter Jurkat cells for 24h at 37°C. CD16 activation was then measured by firefly Luciferase using Bright-Glo (Promega) and an Enspire plate reader (Perkin-Elmer). Luminescence signal of each well was normalized over the luminescence measured for the coculture of 293T-S-D614G and Jurkat CD16 NFAT Luc in the absence of sera.

### S-fuse neutralization assay

The day before the assay, 16.10^3^ S-Fuse cells (8.10^3^ U2OS-ACE2-GFP1-10 & 8.10^3^ U2OS-ACE2-GFP11) were plated in μClear black 96-well plate. The next day, the virus was incubated with serum or antibody dilutions in DMEM 10%FCS 1%PS for 15min and was added to the cells. Plates were incubated for 18h and fixed with PFA 4%. Plates were then washed and stained with Hoechst 33342. Images were acquired using the Opera Phenix High-Content Screening System (PerkinElmer). GFP area, number GFP and DAPI objects per well were determined using Harmony 191 High-Content Imaging and Analysis Software (PerkinElmer, HH17000012, v.5.0).

### Flow virometry

Virus suspensions were stained with Syto24 (ThermoFisher S7559, 20µM final) and indicated antibodies (1 µg/ml diluted in PBS) for 15min at room temperature. Next, a goat anti-human IgG AlexaFluor 647 (1µg/ml final) was added to the mix and incubated for another 15min. Finally, an equal volume of PFA 4% was added for 15min before further diluting sample in 0.02µm-filtered PBS and analyzed with the Cytoflex Nano (Beckman Coulter) and FlowJo.

### Statistical analysis

Graphpad Prism V10.6.1 was used for statistical analysis.

## Results

### TMPRSS2 impairs antibody binding to membrane-bound spike

To investigate the impact of TMPRSS2 on the binding of anti-S mAbs, we transfected 293T cells stably expressing the D614G S protein (293T-S-D614G) with TMPRSS2-mScarlet-I (**Fig. 1A**) and stained them with either Cv2.9116 (pan-coronavirus anti-S2) or S309 (anti-RBD; parental version of sotrovimab). Flow cytometry analysis revealed that TMPRSS2 expression reduced significantly the median fluorescence intensity (MFI) of both mAbs by 73% and 40%, respectively (respective ranges 74,7%-81,4% and 40,5%-45%, **Fig. 1B** and **1C**). The catalytically inactive TMPRSS2 mutant S441A (TMPRSS2 S441A) does not reduce mAbs binding (**Fig. 1B** and **1C**). Treatment of TMPRSS2-transfected 293T-S-D614G with the TMPRSS2 inhibitor camostat resulted in an increase of the MFI of both Cv2.9116 and S309 (**Fig. S1A**). In the same conditions, the MFI of an anti-MHC-I mAb (clone W6/32) is not decreased by TMPRSS2 (**Fig. S1B**).

**Figure 1.**
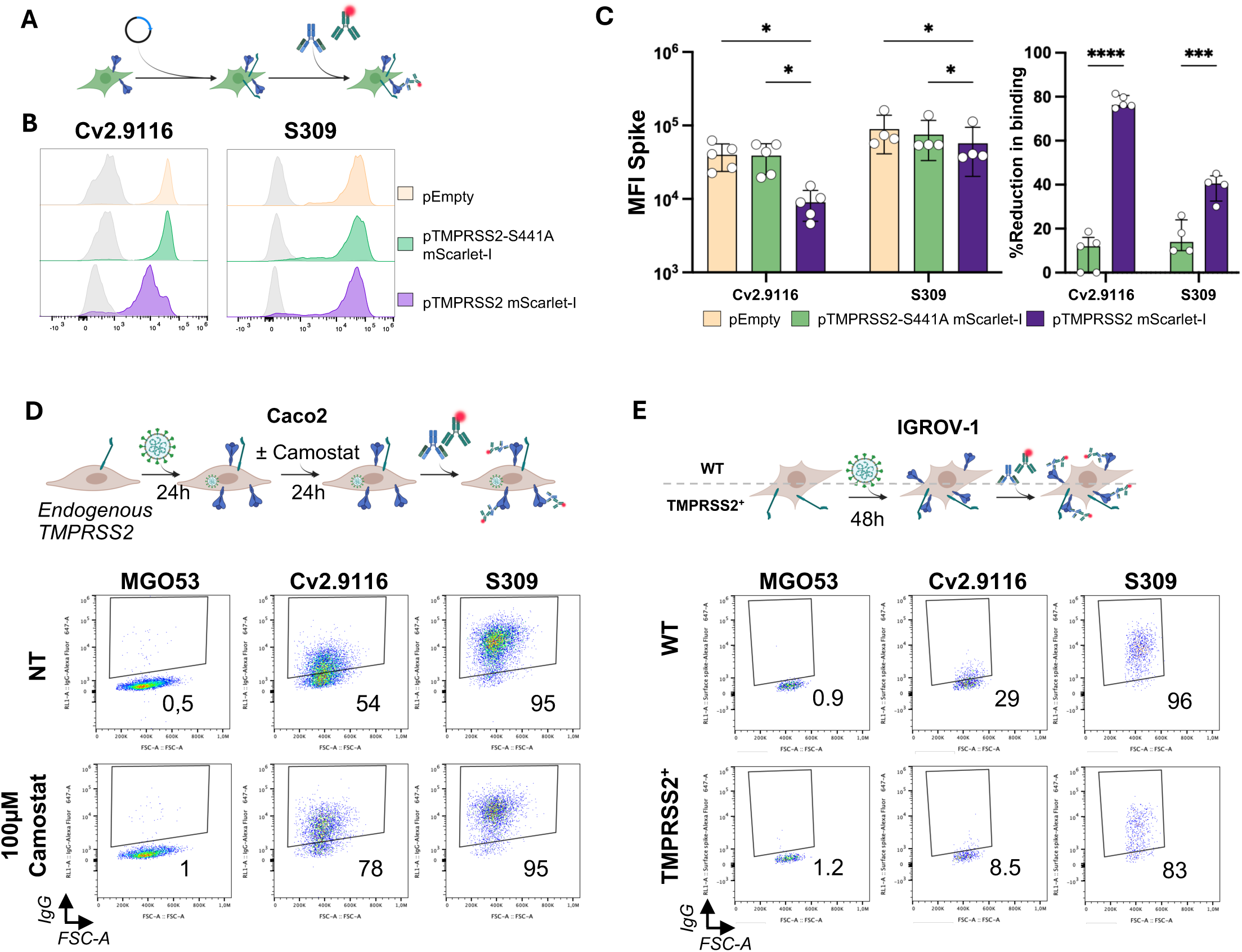
TMPRSS2 decreases mAbs binding to cell surface Spike. **(A**) Experimental procedure. 293T-S-D614G were transfected with TMPRSS2-mScarlet-I plasmid overnight before surface staining with either Cv2.9116 or S309. **(B)** Representative histogram of anti-S binding for each condition. Grey line represents MGO53 (isotype) control for each transfection condition (colored) for Cv2.9116 and S309. **(C)** Raw MFI and %reduction in S binding for Cv2.9116 and S309 for each transfection condition. Median of >3 independent experiments are represented with 95% Confidence Intervals (CIs). **(D)** Experimental procedure for Caco2 cells infection and representative dot plots of anti-S IgG quantification in NCP positive cells (out of 3 independent experiments). **(E)** Experimental procedure for IGROV-1 cells infection and representative dot plots of anti-S IgG quantification in NCP positive cells (out of 3 independent experiments).

Then, we generated Caco2-S-D614G cells and treated them with 1000 µM camostat (a broad serine-proteases inhibitor) 24h before staining with Cv2.9116 and S309 (**Fig. S1C**). The ability of both mAbs to bind to Caco2-S-D614G was increased by camostat from 31 to 34% and from 76 to 81% for Cv2.9116 and S309, respectively. We further infected Caco2 cells with SARS-CoV-2 (variants D614G and XBB.1.5) for 24h before adding Camostat and quantified the binding of anti-spike Cv2.9116 and S309 (**Fig. 1D, S1E** and **Fig. S1D**). Camostat treatment increased the binding of Cv2.9116 to infected cells from 54% to 78% and from 8,2% to 25%, on D614G and XBB.1.5-infected cells, respectively. S309 binding was increased only against XBB.1.5 (from 76% to 81%; Fig. **S1E**). We also infected IGROV-1 cells overexpressing TMPRSS2 with D614G and XBB.1.5 variants and observed a reduction in Cv2.9116 and S309 binding as compared to WT cells (**Fig. 1E, Fig. S1F** and **S1G**).

Finally, we used flow cytometry to compare TMPRSS2 levels across the different cellular systems, including the gold standard human nasal epithelial cells differentiated at the air-liquid interface (hNEC-ALI) model, which express physiological levels of TMPRSS2 (**Fig. S1H**). Transfected 293T cells express the highest levels of TMPRSS2 and Caco2 cells express the lowest and hNEC-ALI cells express intermediate levels (**Fig. S1H**). Overall, our data show that TMPRSS2 decreases the binding of anti-S mAbs to S-expressing and infected cells. This effect relies on the catalytic activity of TMPRSS2 and is specific to S.

### Anti-S2 antibodies exhibit an increased sensitivity to TMPRSS2-mediated evasion

We next investigated whether TMPRSS2 decreases the binding of different mAbs equally. To this end, we used a panel of 39 mAbs recognizing different region on S ectodomain **(Table 1)**. Over-expression of TMPRSS2 in 293T-S-D614G (**Fig. 2A**) and 293T-S-BA.5 (**Fig. S2A**) cells decreases the binding of all tested mAbs. Anti-RBD mAbs show an average 36,1±2,6% decrease in binding upon TMPRSS2 expression, regardless of their RBD class **(Fig. 2B and 2C)**. Among anti-S2 mAbs, TMPRSS2 expression decreases the binding of anti-FP and non-FP mAbs by 46±7% and 72,9±3,5%, respectively. Similarly, not all NTD epitopes behave in the same way, with a larger binding decrease observed for supersite rather than non-supersite mAbs (55,3±12,5% vs 31,6±3,1%). The reduction in binding upon TMPRSS2 expression negatively correlates with binding intensity in the absence of TMPRSS2 **(Fig. 2D)**. This analysis highlights the epitope specificity of TMPRSS2-mediated antibody evasion, as the group that is strongly impacted is composed of anti-S2 (both FP and non-FP) and anti-NTD non-supersite mAbs. In parallel, the group composed of anti-NTD supersite and all anti-RBDs mAbs is less affected. These results align and expand our previous observations that Cv2.9116 binding to infected cells is more affected by TMPRSS2 than S309. Overall, mAbs of distinct specificities are not equally sensitive to TMPRSS2 overexpression, with those targeting the S2 subunit being the most impacted.

**Figure 2.**
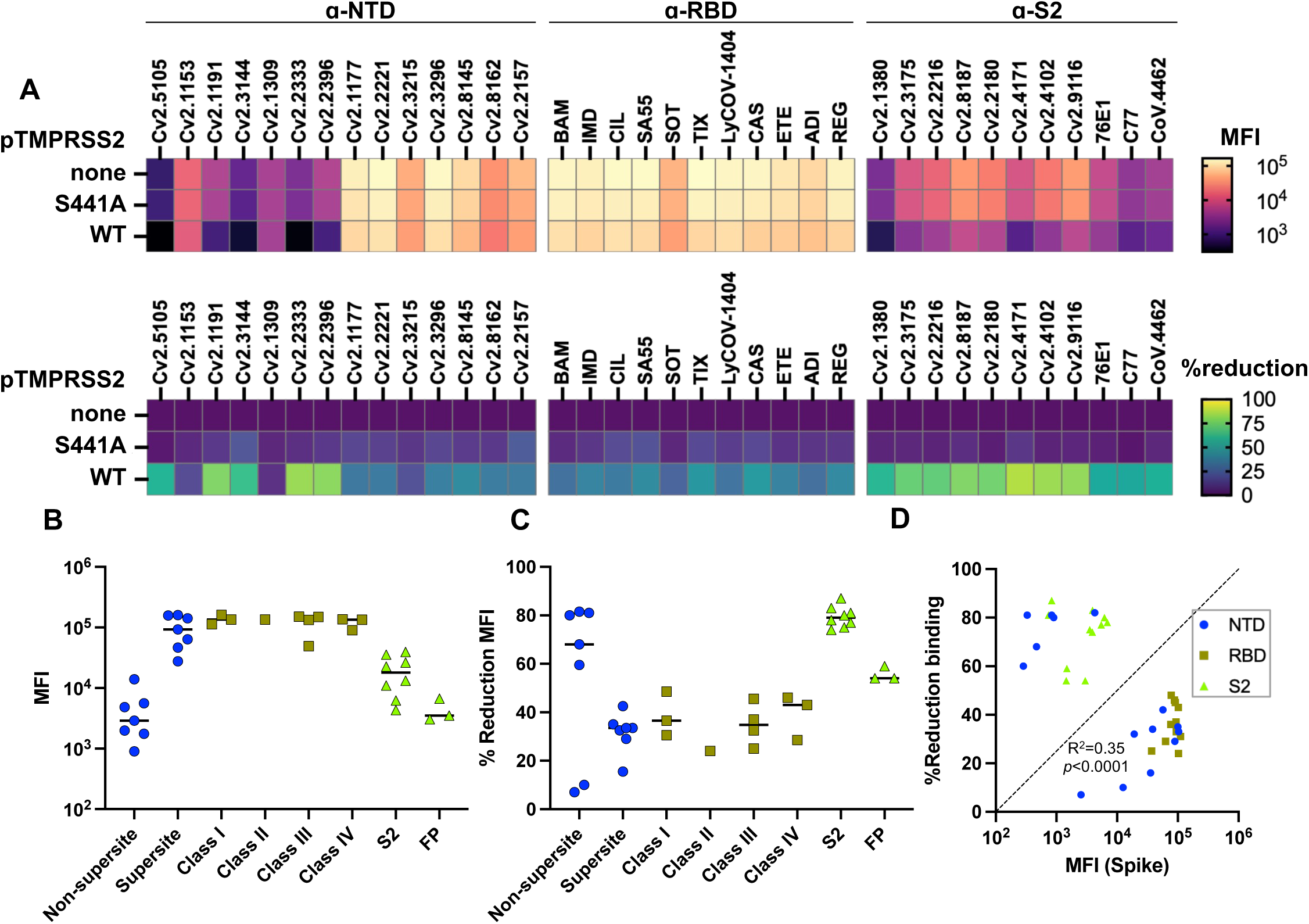
TMPRSS2-mediated decrease in mAb binding varies across Spike epitopes. **(A**) Raw MFI of each mAb and subsequent reduction in binding after TMPRSS2 transfection in 293T-S-D614G are clustered by epitope specificity. Each column is a mAb and each square represents the median value from at least 3 independent experiments. **(B)** Raw MFI and **(C)** reduction in binding to 293T-S-D614G are represented by mAb class and epitope. **(D)** Correlation between the reduction in binding upon TMPRSS2 expression and MFI of binding in the absence of TMPRSS2 for each mAb. A spearman regression was performed.

### TMPRSS2 expression reduces antibody recognition of SARS-CoV-2 variants and other members of the Coronaviridae family

We further aimed to evaluate the conservation of TMPRSS2-mediated decrease of S recognition by mAbs across SARS-CoV-2 variants. We co-transfected TMPRSS2 and S proteins from 10 different SARS-CoV-2 variants spanning the pandemic (from D614G to KP.3.1.1) and evaluated Cv2.9116 binding **(Fig. 3A)**. The binding consistently decreases with TMPRSS2 expression, independently of the viral variants **(Fig. 3B** and **S3A)**. Using the same methodology, we investigate Cv2.9116 binding to different coronavirus S proteins including human coronavirus 229E (alphacoronavirus) and SARS-CoV-2-related bat coronaviruses (RaTG13 and BANAL 20-236). Of note, OC43 could not be included as its S protein is not exported at the plasma membrane (Sadasivan et al., 2017). TMPRSS2 significantly lowers Cv2.9116 binding to 229E and RaTG13 S, whereas binding to BANAL 20-236 S is not affected in any condition **(Fig. 3C**, **Fig. S3B** and **S3C)**. Note that the low levels of NL63-S expression impair accurate quantification, but we still observe that catalytically active TMPRSS2 reduces the binding of Cv2.9116 **(Fig. S3C)**. These results show that the ability of TMPRSS2 to promote the escape of S-expressing cells from anti-S2 antibodies is not restricted to SARS-CoV-2.

**Figure 3.**
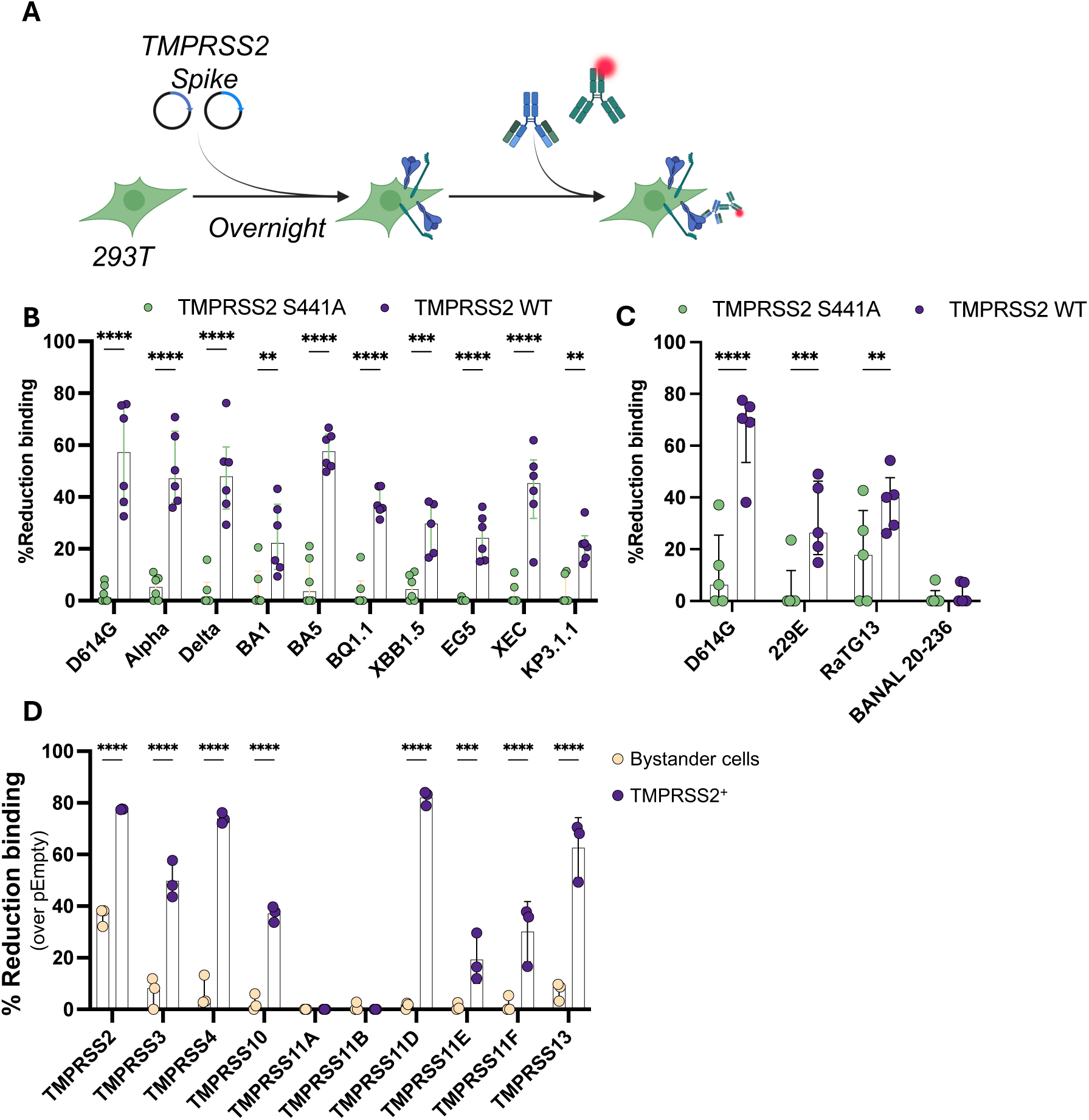
TMPRSS2 reduction of antibody binding is conserved across betacoronaviruses and observed with other TMPRSS2-related proteases. **(A)** Experimental procedure. **(B)** Reduction in binding to SARS-CoV-2 S-positive cells upon TMPRSS2 expression as compared to pEmpty using S from different SARS-CoV-2 variants. **(C)** Reduction in binding to S-positive cells upon TMPRSS2 as compared to pEmpty using S plasmids of different betacoronaviruses. **(D)** Reduction in binding of Cv2.9116 to 293T-S-D614G transfected with different TMPRSS. Median with 95% CI are represented and each point represents a single biological replicate. Statistical analyses were performed using two-sided ANOVA, *p*<0.05 = *, *p*<0.01 = **, *p*<0.001 = ***, *p*<0.0001 = ****.

### Several members of the TMPRSS family decrease antibody binding on SARS-CoV-2 S

Next, we evaluated the capacity of 8 proteases from the TMPRSS family to decrease antibody recognition of SARS-CoV-2 S. For this, we transfected myc-conjugated TMPRSS proteases into 293T-S-D614G cells and assessed Cv2.9116 binding **(Fig. S3D** and **S3E)**. TMPRSS11A and 11B slightly increase Cv2.9116 binding to 293T-S-D614G in TMPRSS-expressing cells as compared to bystander cells **(Fig. S3E).** All other TMPRSS proteases induce significant reductions in Cv2.9116 binding to 293T-S-D614G **(Fig. 3D)**. Interestingly, TMPRSS2 transfection reduces binding of Cv2.9116 on both transfected and bystander cells, suggesting a *cis-* and *trans*-action on S for TMPRSS2. Altogether, this data shows that the capacity to decrease antibody recognition of SARS-CoV-2 S is shared by distinct members of the TMPRSS family.

### TMPRSS2 promotes S conformational rearrangements and increases S1 shedding

We sought to investigate the mechanism by which TMPRSS2 decreases mAbs binding to 293T-S-D614G. To this aim, we employed mAbs targeting a linear epitope within the HR2 region of S2 (Cv2.3132; (Planchais et al., 2024)) and a quaternary epitopes of the RBD (2-43; (Liu et al., 2020)) to determine whether S is impacted in either quantity or conformation, respectively. As a control of TMPRSS2 activity we used Cv2.9116 **(Fig. 4A, 4B** and **4C)**. As described above, overexpression of TMPRSS2 decreases Cv2.9116 staining by a median of 83,3%. In contrast, the binding of Cv2.3132 is reduced by only a median of 15%, demonstrating that a change in overall S quantity cannot fully explain the observed decrease in Cv2.9116 binding. 2-43 is decreased by a median 49,1%, consistent with either shedding of S1, a major conformational change of S, or both **(Fig.4A, 4B** and **4C)**. We then perform an ACE2 binding assay using a soluble ACE2 ectodomain (**Fig. 4D**), which shows a reduction in the MFI upon TMPRSS2 transfection from 4266 to 1622. Therefore, we measured soluble S1 in the supernatant of 293T-S-D614G cells either overexpressing TMPRSS2 or not by ELISA **(Fig. 4E)**. Catalytically active TMPRSS2 increases the levels of soluble S1 by an average of 1.8-fold over both empty control plasmid and TMPRSS2 S441A (**Fig. 4E**). These data indicate that TMPRSS2 elicits S1 shedding. Note that we observe a similar shedding of S1 upon TMPRSS2 transfection with 293T-S-BA.5 (**Fig. S4A**, **B**, **C** and **D**). Taken together, these results demonstrate that TMPRSS2 marginally affects S surface levels, but rather promotes conformational changes characterized by S1 shedding and a decreased susceptibility to anti-S2 antibodies.

**Figure 4.**
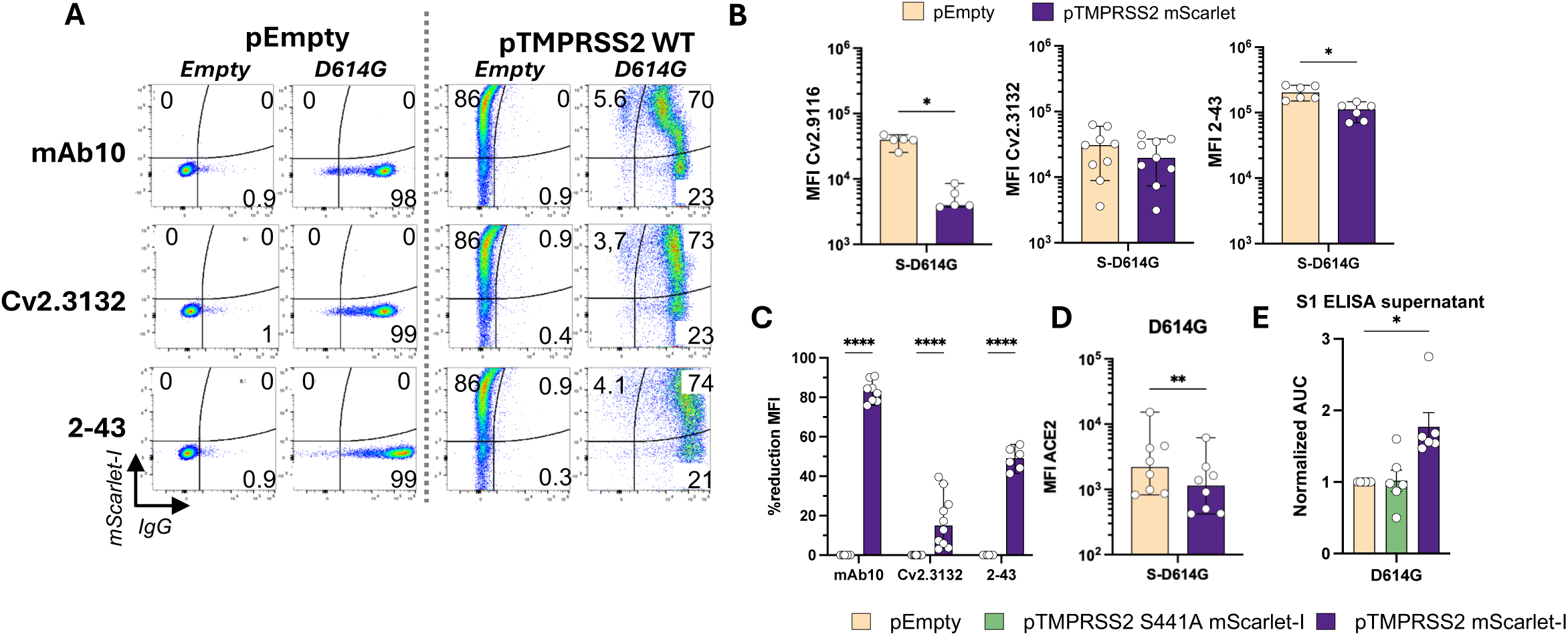
TMPRSS2 effect on antibody binding is mediated by conformational changes of Spike. **(A**) Dot plot of Cv2.9116, Cv2.3132 and 2-43 binding to 293T-empty or 293T-S-D614G upon transfection of pEmpty or TMPRSS2 WT. **(B)** Quantification of Cv2.3132, 2-43 and Cv2.9116 binding to 293T-S-D614G after TMPRSS2 transfection and **(C)** corresponding reduction in binding. Mann-Whitney analysis was performed, *p*<0.05 = * for MFI and reduction in MFI was analyzed using two-sided ANOVA, *p*<0.0001 = ****. **(D)** Soluble ACE2 MFI to 293T-S-D614G. **(E)** Normalized Area Under the Curve (AUC) quantified by S1 ELISA in supernatants after overnight transfection in 293T-S-D614G. Each point is a biological replicate and median and 95% CI are represented. Mann-Whitney analysis was performed, *p*<0.05 = *.

Considering that we and others have previously reported that TMPRSS2 increases S fusogenicity (Bestle et al., 2020; Buchrieser et al., 2020), TMPRSS2-mediated S1 shedding and lower ACE2 binding appears counter-intuitive. Therefore, we evaluated cell-cell fusion in the same conditions in which we observed a decrease of mAbs binding by TMPRSS2, using an acceptor-donor GFP-split assay (**Fig. 5A**). Regardless of whether TMPRSS2 is expressed by donor, acceptor or both cells, it increases S fusogenicity with ACE2-expressing (**Fig. 5B**, **5C**, and **S4E**). Interestingly, TMPRSS2 does not increase ACE2-independent fusion, which remains comparable to background (**Fig. 5D, 5E**, and **S4E**), suggesting that the decrease in ACE2 binding is not compensated by an unspecific triggering of fusion. Similarly, TMPRSS2 increases only ACE2-dependent cell-cell fusion of 293T-S-BA.5 (**Fig. S4F** and **S4G**). Overall, these results suggest that while TMPRSS2 reduces the number of S capable of binding ACE2, the remaining S are more efficient at eliciting fusion upon ACE2 engagement.

**Figure 5.**
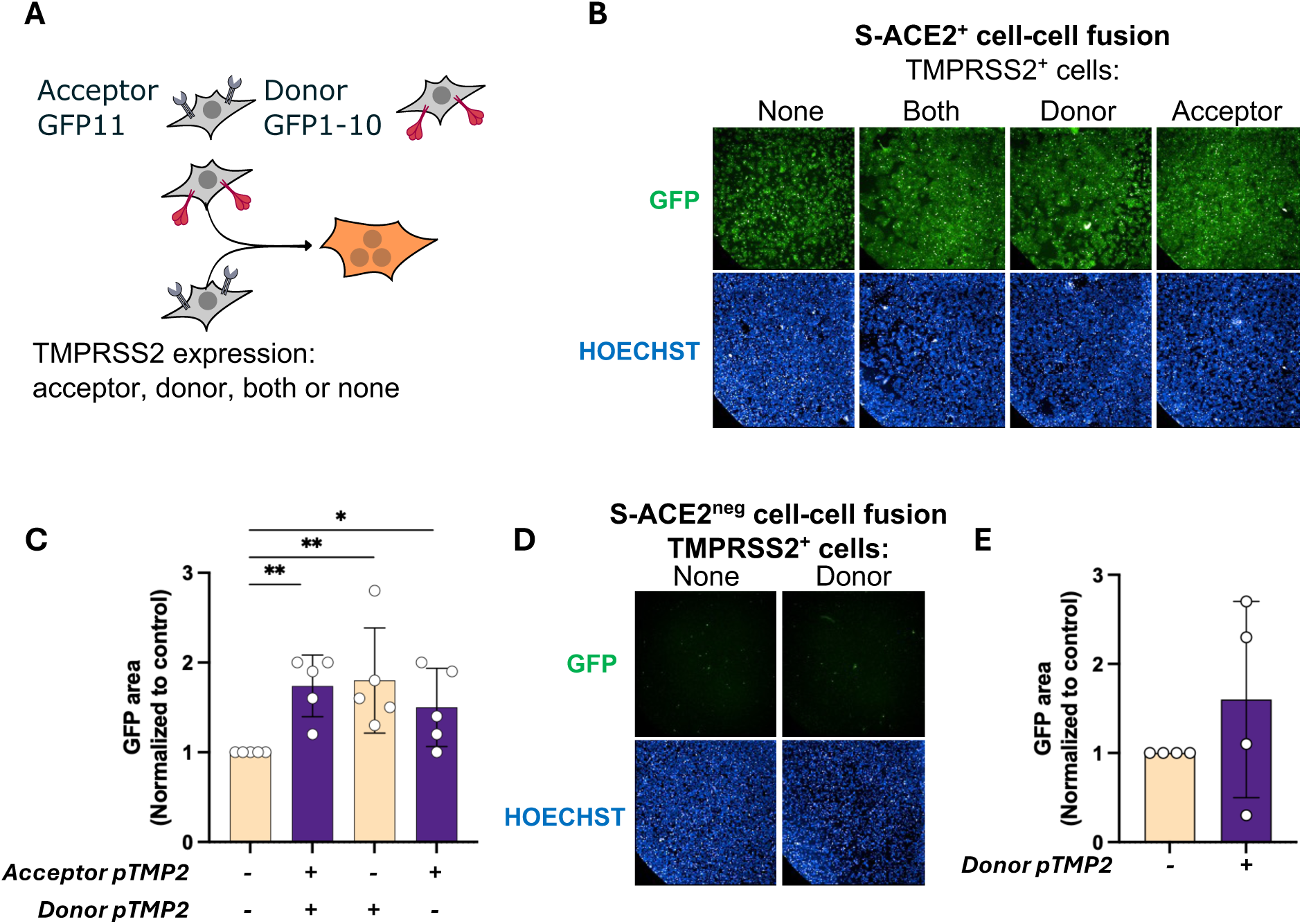
Evaluation of TMPRSS2 on S-mediated syncytia formation. **(A)** Experimental design. **(B)** Representative images of spike-mediated fusion with ACE2^+^ overexpressing 293T after TMPRSS2 transfection and **(C)** corresponding quantification. Statistical analysis was performed using multiple Mann-Whitney tests, *p*<0.05 = *, *p*<0.01 = **. **(D)** Representative images of spike-mediated fusion with WT overexpressing 293T after TMPRSS2 transfection and **(E)** corresponding quantification. Shown are median and 95% CI of all experiments while each point represents individual values from biological replicates.

### TMPRSS2 decreases serum-mediated ADCC of S-expressing cells

Next, we sought to investigate whether TMPRSS2 expression decreases the binding of polyclonal anti-S antibodies. We used immune sera from individuals with hybrid immunity (n = 15; Wuhan infection, followed by 3 doses of mRNA vaccination; **Table 2**). Transfection of TMPRSS2 reduces the binding of these sera to 293T-S-D614G by 52% (min 45,5 - max 56,3), as measured by IgG staining (**Fig. 6A**). Similarly, treatment of Caco2-S-D614G with Camostat increases the binding of polyclonal IgG and IgA antibodies (**Fig. S5A**). As expected, TMPRSS2 expression does not change the binding of pre-pandemic sera, which remained negative (**Fig. S5B**). Infected Caco2 cells with D614G or XBB1.5 variants show a similar increase in serum IgG binding to infected cells after 24h of Camostat treatment **(Fig. 6B** and **S5C)**. In IGROV-1 cells, overexpression of TMPRSS2 lowers IgG binding to cells infected with either D614G or XBB1.5 **(Fig. 6C** and **S5D)**. This data indicates that endogenous levels of TMPRSS2 and/or serine-proteases decrease the recognition of SARS-CoV-2 infected cells by antibodies present in the sera of hybrid immune individuals.

**Figure 6.**
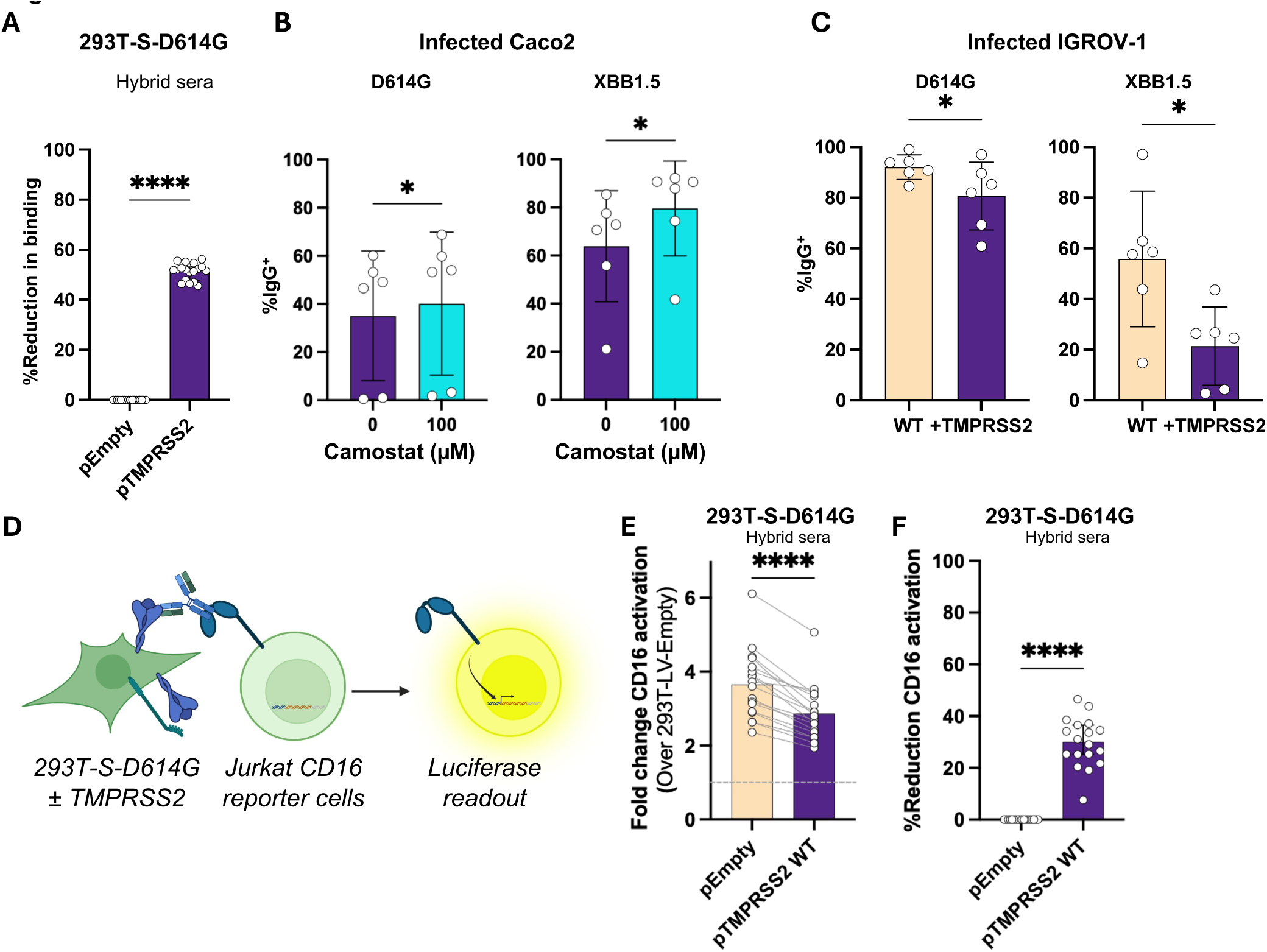
Seropositive serum binding evaluation in TMPRSS2 context. **(A)** Reduction of anti-S Abs binding from serum of individuals with hybrid immunity to 293T-S-D614G expressing or not TMPRSS2 **(B)** Percentage of IgG bound to D61G4- or XBB1.5-infected Caco2 cells by anti-S from the same sera after overnight treatment with camostat. **(C)** Percentage of IgG bound to D61G4 or XBB1.5 infected IGROV-1 cells expressing or not TMPRSS2 by the same sera. **(D)** Experimental procedure for CD16 reporter coculture assay and **(E)** corresponding quantification of CD16 activation. Displayed are the median and 95% confidence interval. Points represent individual values from biological replicates. Wilcoxon statistical analysis was performed, *p*<0.05 = *, *p*<0.0001 = ****

To evaluate if lower binding of serum IgGs from individuals with hybrid immunity translates into lower ADCC induction, we co-cultivate TMPRSS2-transfected 293T-S-D614G with CD16-NFAT reporter Jurkat cells (**Fig. 6D**). We observe that TMPRSS2 transfection significantly lowers CD16 activation median from 3,7 to 2,8, representing a median 30% (min 38,9% - max 46,5%) decrease (**Fig. 6E**). This highlights that the capacity of TMPRSS2 to lower antibody binding results in impaired CD16 engagement and downstream signaling, which is an accurate indicator of ADCC induction against infected cells (Dufloo et al., 2021).

### TMPRSS2 expression in infected cells marginally affects viral progeny

Finally, we aim to determine whether TMPRSS2 expression in infected cells can impact the antibody response to the viral progeny. For this, we produced two independent batches of the D614G variant in IGROV-1 cells either expressing TMPRSS2 or not to analyze the released viruses for infectivity and antibody sensitivity (**Fig. 7A**). We observed that TMPRSS2 expression in infected cells accounts for higher infectious titers at 2 days post-infection (3,4.10^6^ vs 1.10^6^), but the difference diminishes at 3 dpi (**Fig. 7B**), possibly because of an increased cytopathic effect in TMPRSS2-expressing cells (not shown). Interestingly, viral genome copies in the supernatant are not significantly different between cellular backgrounds. Thus, the ratio of infectious genomes (RNA copies per infectious units) is slightly higher for viruses produced in IGROV-1 TMPRSS2^+^ at 2 dpi. These results indicate that TMPRSS2 presence in infected cells renders viral progeny slightly more infectious at early time points.

**Figure 7.**
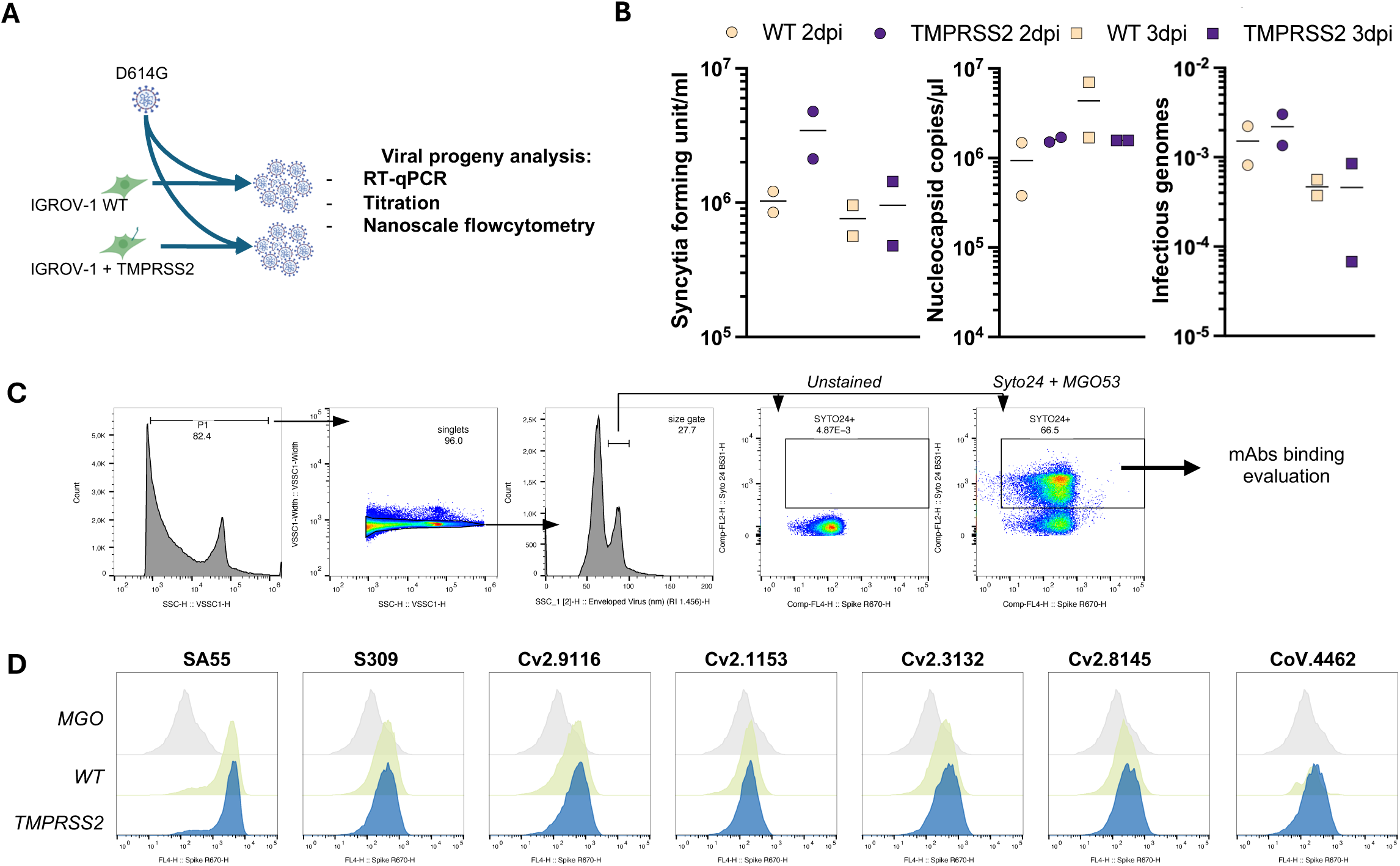
Evaluation of TMPRSS2 status in infected cells on the viral progeny. **(A)** Schematic outline of the procedure. **(B)** S-Fuse titration, nucleocaspid RT-qPCR and Infectious genomes of supernatant from two independently produced stocks of D614G viruses, measured in duplicates. **(C)** Flow virometry gating strategy to analyze anti-S binding to viral population from WT IGROV-1 cells. **(D)** Representative histograms of binding intensity. WT and TMPRSS2 conditions are represented in green and blue, respectively.

We further employ flow virometry to evaluate antibody binding to viral particles produced in each cellular background, using a selection of mAbs (SA55, S309, Cv2.9116, Cv2.3132, Cv2.8145 and CoV44-62). We identify viral particles as events of around 80-90 nm in size and positive for the nucleic acid dye Syto24 (**Fig. 7C**). We do not observe any difference in mAb binding (**Fig. 7D** and **S6**), apart from SA55, which exhibits a modest decrease. Altogether, these results suggest that TMPRSS2 expression in infected cells has only moderate effects on S antigenicity in the viral progeny.

## Discussion

Our study shows that TMPRSS2 makes the plasma membrane-associated S less sensitive to recognition by antibodies and subsequent ADCC. We show that this activity requires a catalytically active TMPRSS2. Its presence only marginally affects surface S level at the membrane, indicating that TMPRSS2 does not reduces antibody binding because of a degradation of S. We observe an increased S1 shedding, which likely explains the decrease in binding of anti-S1 antibodies (i.e. anti-RBD and anti-NTD). Anti-S2 antibodies are more affected by TMPRSS2-mediated evasion, a finding that cannot be explained by the shedding of S1. Therefore, the S2 domain must adopt a conformation less susceptible to antibody binding. Recent studies have identified an intermediate state of the S protein, characterized by partial shedding and reversible rearrangement of S2 (Olmedillas et al., 2025). Combining our data with these structural insights, we propose a model where TMPRSS2 destabilizes S through cleavage, enabling S to adopt a conformation in-between pre- and post-fusion states, which is less recognized by antibodies.

TMPRSS2 cleaves S at multiple sites (Fraser et al., 2022). We do not identify the contribution of each of those sites to the phenotype. At least two of these cleavage sites seem relevant to the phenomenon reported here. First, TMPRSS2 can cleave the Arg-Ser peptide bond of the furin cleavage site (R–X–[K/R]–R↓–S) and therefore promotes shedding by favoring S1 and S2 dissociation. Second, cleavage of the S2’ site can account for the preferential impact on anti-S2 antibodies by disrupting local epitopes required for anti-S2 binding, as many FP epitopes overlap with the S2’ site (Dacon et al., 2022; Low et al., 2022; Rosen et al., 2024). It is also plausible that cleavage of S2’ increases the overall flexibly of S2 (Olmedillas et al., 2025), enabling this subunit to sample conformations that would hide key epitopes. Cleavage of S2’ can occur on the closed trimer independently of S1 shedding (Fraser et al., 2022), as the Arg is not buried within the trimer, explaining why TMPRSS2 can reduce antibody binding to S in the absence of ACE2. Our study clearly demonstrates the overall impact of TMPRSS2 on S recognition by antibodies, but how each cleavage sites combine to affect overall S antigenicity remains to be elucidated.

Our results also suggest that TMPRSS2 affects S present at the plasma membrane of infected cells more than S proteins incorporated into viral particles. This finding aligns with SARS-CoV-2 budding in the ERGIC compartment and exiting cells through a mixed lysosomal-secretion pathway (Chen et al., 2021; Miura et al., 2023; Scherer et al., 2022). This intracellular assembly of viral progeny most likely prevents contacts between virion-associated S and activated TMPRSS2, which is preferentially located at the plasma membrane (Zhang et al., 2022). Importantly, shedding of S at the plasma membrane may still interfere with the neutralization of viral particles by generating soluble S1 decoy, as previously shown for SARS-CoV-1 (Glowacka et al., 2011). A similar mechanism can operate in SARS-CoV-2 infected cells.

As the physiological antibody response to SARS-CoV-2 infection and vaccination is inherently polyclonal, we used immune positive sera from a hybrid immunity cohort. TMPRSS2 decreases S binding and ADCC potency of these serum antibodies. Although the CD16 reporter assay used differs from genuine ADCC assays, we have previously reported a strong correlation between this assay and NK cell-mediated ADCC of SARS-CoV-2 infected cells (Dufloo et al., 2021).

SARS-CoV-2 variants have evolved to escape anti-S binding and neutralization (Planas et al., 2024, 2022, 2021; Uraki et al., 2026). Viral variants also differ in the proteolytic processing of S, as the efficiency of the S1/S2 cleavage gradually increased while the usage of TMPRSS2 decreased with Omicron emergence. TMPRSS2 decreases antibody binding against all variants tested, indicating that it does not correlate with TMPRSS2 usage bias (Meng et al., 2022). Differential TMPRSS2 impact on the anti-S2 pan-coronavirus Cv2.9116 binding across the *Coronaviridae* family is also informative, as all coronaviruses share a conserved S2’ cleavage sequence, although the non-SARS-CoV-2 viruses tested here lack the FCS (Chan and Zhan, 2021; Hoffmann et al., 2020; Temmam et al., 2022). As previous studies reported multiple cleavage sites by TMPRSS2 in SARS-CoV and SARS-CoV-2 S (Fraser et al., 2022; Glowacka et al., 2011), it is plausible that other cleavage sites may be involved, and possibly differ across *Coronavidae*.

It is well established that anti-S2 Abs are cross-reactive, elicit effector functions, mediate *in vivo* protection in animal models and protects against severe disease in clinical studies (Dacon et al., 2022; Guenthoer et al., 2024; Ng et al., 2022, 2021; Stravalaci et al., 2022; Zhou et al., 2023). Hence, lowering the binding of anti-S2 antibodies by TMPRSS2 could be a mechanism by which SARS-CoV-2 escapes these cross-reactive antibodies.

We also show that other TMPRSS mediate Ab-escape. This aligns with previous reports indicating that TMPRSS4, 11D, 11E and 13 increase SARS-CoV-2 infection *in vitro* (Kishimoto et al., 2021; Zang et al., 2020). TMPRSS2 expression level in dissociated primary ciliated cells is similar to that observed in transfected 293T. However, inhibition of TMPRSS2 (as well as other TMPRSS proteases) with Camostat mesylate treatment showed no significant efficacy against COVID-19 in clinical trials (Khan et al., 2024). Given our results, it would be interesting to investigate the possible antiviral synergy between Camostat and antibodies, especially the ones targeting S2. Overall, future work will uncover how cellular proteases cooperate to modulate S conformation and antibody escape to reveal potential new therapeutic targets.

In conclusion, our results highlight the diverse effects of TMPRSS2 on the SARS-CoV-2 cycle and uncover a new mechanism by which cells infected by SARS-CoV-2 avoid the humoral immune response.

## Acknowledgments

We thank members of the Virus and Immunity Unit for discussions and help in the preparation of this manuscript. We thank Jocelyne Creff and Fabienne Tzvetkov-Ricard for their technical and administrative assistance. We thank N. Aulner, N. Mahtal, J. Fernandes and the UtechS Photonic BioImaging core facility (Institut Pasteur). We thank Divya Unni and the Institut Pasteur grant office for their invaluable assistance in funding acquisition. We thank all the staff members of the Occupational Health and Medicine Department of the Hospices Civils de Lyon and staff members who contributed to the COVID-Ser clinical, all the clinical research associates for excellent work, and all the members of the clinical research and innovation department for reactivity (DRS, Hospices Civils de Lyon), in particular Pr Jean-Baptiste Fassier. Last, we thank all the patients and the health care workers for participation in this clinical study. Human biological samples and associated data were obtained from NeuroBioTec (CRB HCL, Lyon France, Biobank BB-0033-00046).

## Fundings

This work is funded by Institut Pasteur, ANRS-MIE (D25266 IMMUNO-COVID), the Vaccine Research Institute (ANR-10-LABX-77), and the ANR young investigator program (ANR-23-CE15-0039-01 MACOVA). The funders had no role in the design or writing of the article. UtechS Photonic BioImaging is funded by grant no. ANR-10-INSB-04-01, no. ANR-24-INBS-0005 FBI (BIOGEN) and Région Ile-de-France programme DIM1-Health. This study received funding from the Agence Nationale de Recherches sur le Sida et les Hépatites Virales (ANRS-0154) and Fondation des Hospices Civils de Lyon for the COVID-Ser cohort. We acknowledge support from Institut Pasteur for the use of the Opera Phenix Plus microscope.

## Author contributions

Experimental strategy and design: A.C.C., O.S. and T.B.

Laboratory experiments: A.C.C., A.D.C., C.P., F.P., M.K., M.J.G., E.T., A.W., I.S., F.G.B., P.R., I.F., J.B. and T.B.

Viral strains and key reagents and techniques: I.S., F.G.B., F.P., C.P., P.R. and H.M.

Cohort management and clinical research: S.T.A

Funding acquisition: O.S., H.M. and T.B.

Manuscript writing & editing: A.C.C, F.R., H.M., O.S and T.B

## Conflict of interest

All authors declare no conflicts of interest.

## Inclusion and diversity

We support inclusive, diverse, and equitable conduct of research.

**Figure Suppl 1. TMPRSS2 catalytic activity confers lower anti-S binding.**

**(A)** Dot plots of the effect of simultaneous transfection with TMPRSS2 and Camostat treatment of 293T-S-D614G on Cv2.3132, Cv2.9116 and S309 binding. Representative of three biological replicates. **(B)** MFI of anti-MHC-I W6/32 binding to 293T-S-D614G transfected either with pTMPRSS2 S441A or WT. Each point is a biological replicate, median and 95% CI are represented. **(C)** Dot plots of the effect of overnight Camostat treatment of Caco2-S-D614G Cv2.9116 and S309 binding. Representative of three biological replicates. **(D)** Gating strategy for the analysis of Cv2.9116 and S309 binding to infected Caco2, represented for non-treated Caco2 infected with D614G. **(E)** Dot plots of the effect of overnight Camostat treatment at 1dpi XBB1.5 infected Caco2 on Cv2.9116 and S309 binding. Representative of three biological replicates. **(F)** Gating strategy for the analysis of Cv2.9116 and S309 binding to infected IGROV-1, represented for IGROV-1 WT infected with D614G. **(G)** Dot plots of the effect TMPRSS2 in XBB1.5-infected Caco2 on Cv2.9116 and S309 binding representative of three biological replicates. **(H)** Binding of anti-TMPRSS2 nanobody binding to different cells in a representative histogram (left) where grey lines represent the MGO53 staining and the corresponding plot (left).

**Figure Suppl 2. Extension of epitope screening of TMPRSS2-mediated decrease in mAb binding to 293T-S-BA.5**

Raw MFI of each mAb and subsequent reduction in binding after TMPRSS2 transfection in 293T-S-BA.5 are clustered by epitope specificity. Antibodies showing baseline binding to 293T-S-BA.5 under 3 times the value of MGO53 (irrelevant mAb) were not considered for reduction analysis and showed in grey. Each column is a mAb and each square represents the median value from >3 independent experiments.

**Figure Suppl 3. Evaluation of TMPRSS activity on Cv2.9116 binding to different coronaviruses spike.**

**(A** and **B)** Raw MFI of Cv2.9116 binding in co-transfected 293T cells. Dashed line represents mean MGO53 MFI, as background measurement. MFI signal was analyzed using two-sided ANOVA, *p*<0.05 = *, *p*<0.01 = **, *p*<0.001 = ***, *p*<0.0001 = ****. **(C)** Representative dot plots of co-transfected 293T cells with different betacoronaviruses’ S. **(D)** Representative dot plots of Cv2.9116 binding to 293T-S-D614G cells transfected with different TMPRSS and **(E)** Raw MFI. Each point is a biological replicate, median and 95% CI are represented. Dashed line represents mean MGO53 MFI, as background measurement. MFI signal was analyzed using two-sided ANOVA, *p*<0.05 = *, *p*<0.01 = **, *p*<0.001 = ***, *p*<0.0001 = ****.

**Figure Suppl 4. Extension of the mechanistic analysis of TMRPSS2 effect on mAb binding on S-BA.5 and fusion.**

**(A**) Quantification of Cv2.3132, IgG 2-43 and Cv2.9116 binding to 293T-S-BA.5 after TMPRSS2 transfection and **(B)** corresponding reduction in binding. Mann-Whitney analysis was performed, *p*<0.05 = * for MFI while reduction in MFI was analyzed using two-sided ANOVA, *p*<0.0001 = ****. **(C)** Soluble ACE2 MFI to 293T-S-BA.5. Histograms represent the median and 95% CI of all experiments while each point represents individual values form biological replicates. **(D)** Normalized area under the curve (AUC) quantified by S1 ELISA contained in supernatants after overnight transfection in 293T-S-BA.5. Mann-Whitney analysis was performed, *p*<0.05 = *. **(E** and **F)** Raw values of GFP area in µm^2^ for ACE2-dependent and ACE2-independent S-D614G- and S-BA.5 respectively mediated fusion after TMPRSS2 transfection in donor or acceptor cells. **(G)** GFP area normalized over non-transfected GFP area for S-BA.5 donor cells upon TMPRSS2 transfection. Shown are the median of all experiments and 95% CI with each point representing individual values form biological replicates. Statistical analysis was performed using two-way ANOVA, *p*<0.01 = **.

**Figure Suppl 5. Analysis of serum Abs binding to membrane-bound spike.**

**(A)** Percentages of IgG and IgA positive Caco2-S-D614G upon overnight Camostat treatment. Each point represents the mean of two independent measurements for each donor. Statistical analysis was performed using two-sided ANOVA, *p*<0.0001 = ****. (**B**) Raw MFI of pre-pandemic and seropositive sera to 293T-S-D614G transfected with TMPRSS2 or not. Each point represents mean of three independent measurements for each donor. **(C)** Gating strategy for evaluation of anti-S binding from sera binding gated on nucleocapsid-positive Caco2 cells, exemplified for non-treated D614G-infected cells. (**D**) Gating strategy for evaluation of anti-S binding from sera gated in nucleocapsid-positive IGROV-1, exemplified for WT D614G-infected cells.

**Figure Suppl 6. Quantification of each mAb binding D614G produced in TMPRSS2 IGROV1 cells.** Each point is an independent measurement. Dashed line represents the minimal and maximal MGO53 MFI obtained across the independent experiment. Each viral production has been analyzed at least 2 times. Statistical analysis was performed using 2-sided ANOVA *p*<0.001 = ***.

## Notes

### Competing Interest Statement

The authors have declared no competing interest.

### Summary of Updates

To correct a mistake (sentence cropped) in the abstract and add the supplementary file.

