## Supplementary figures and images for "TMPRSS2 reduces antibody recognition of SARS-CoV-2 spike"

### Supplementary Figures

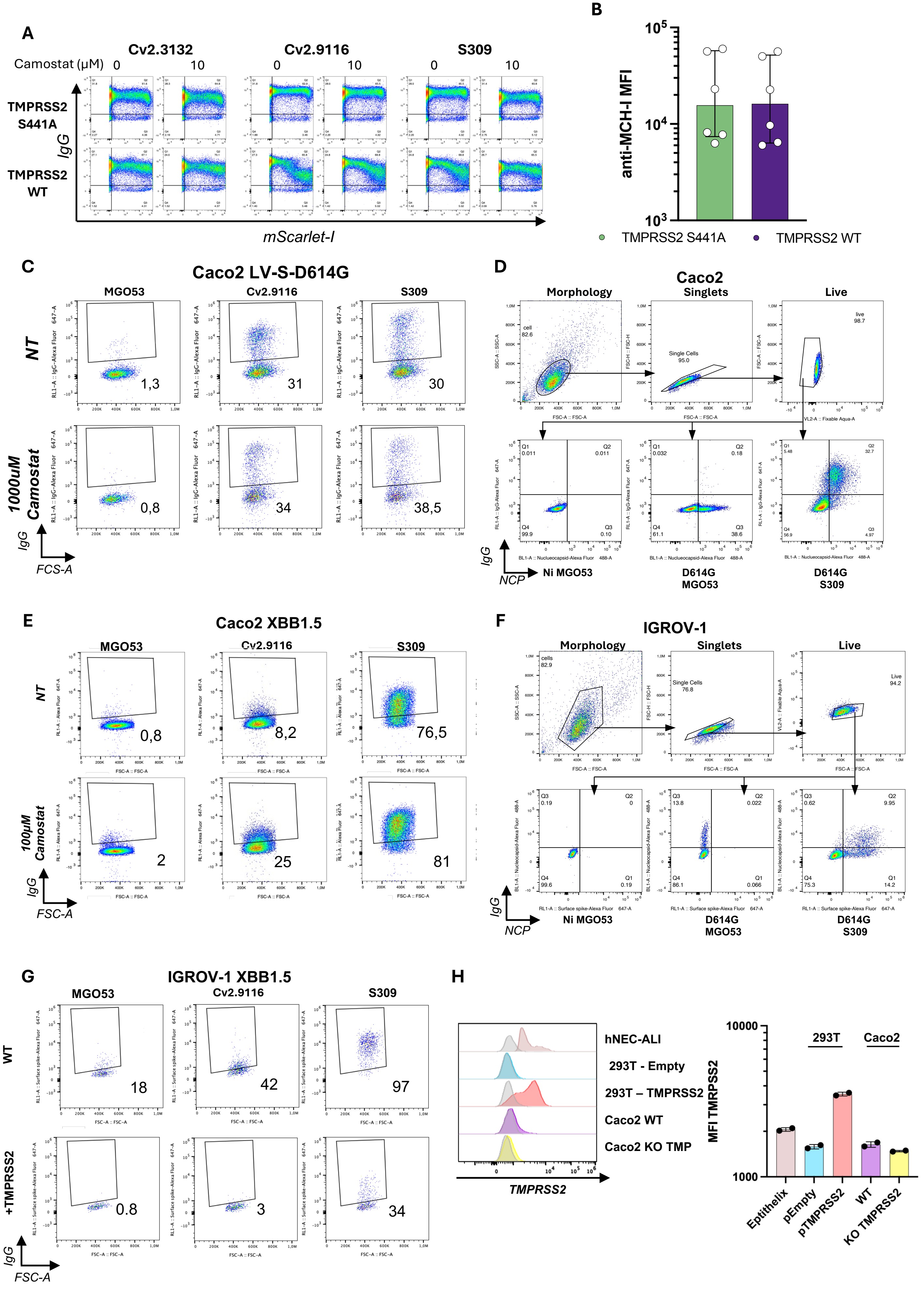

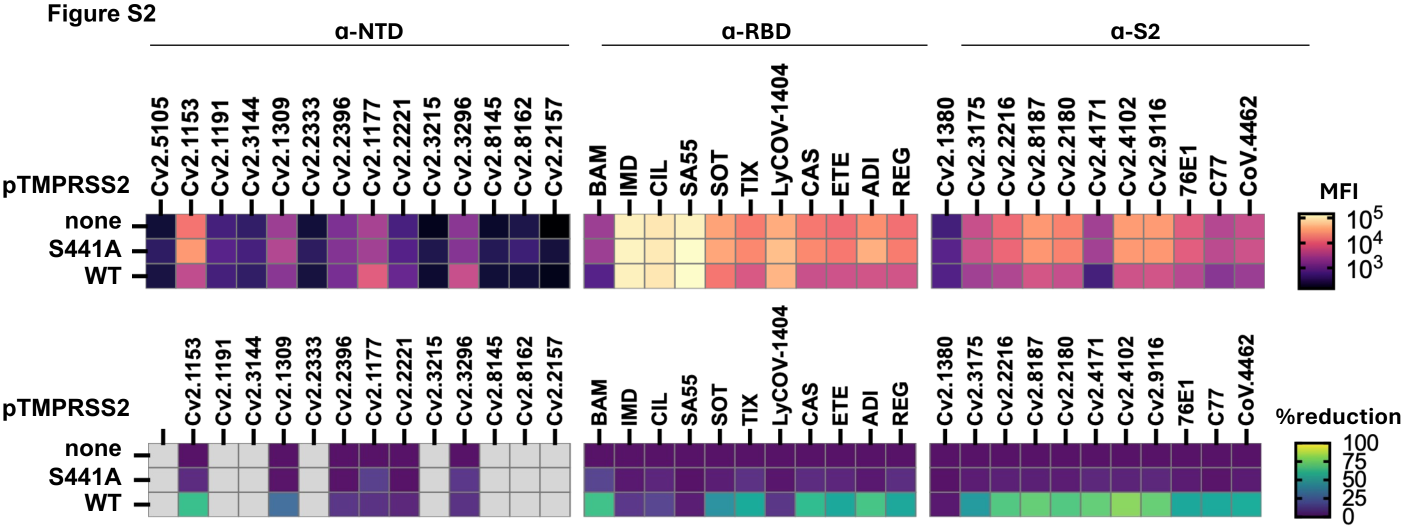

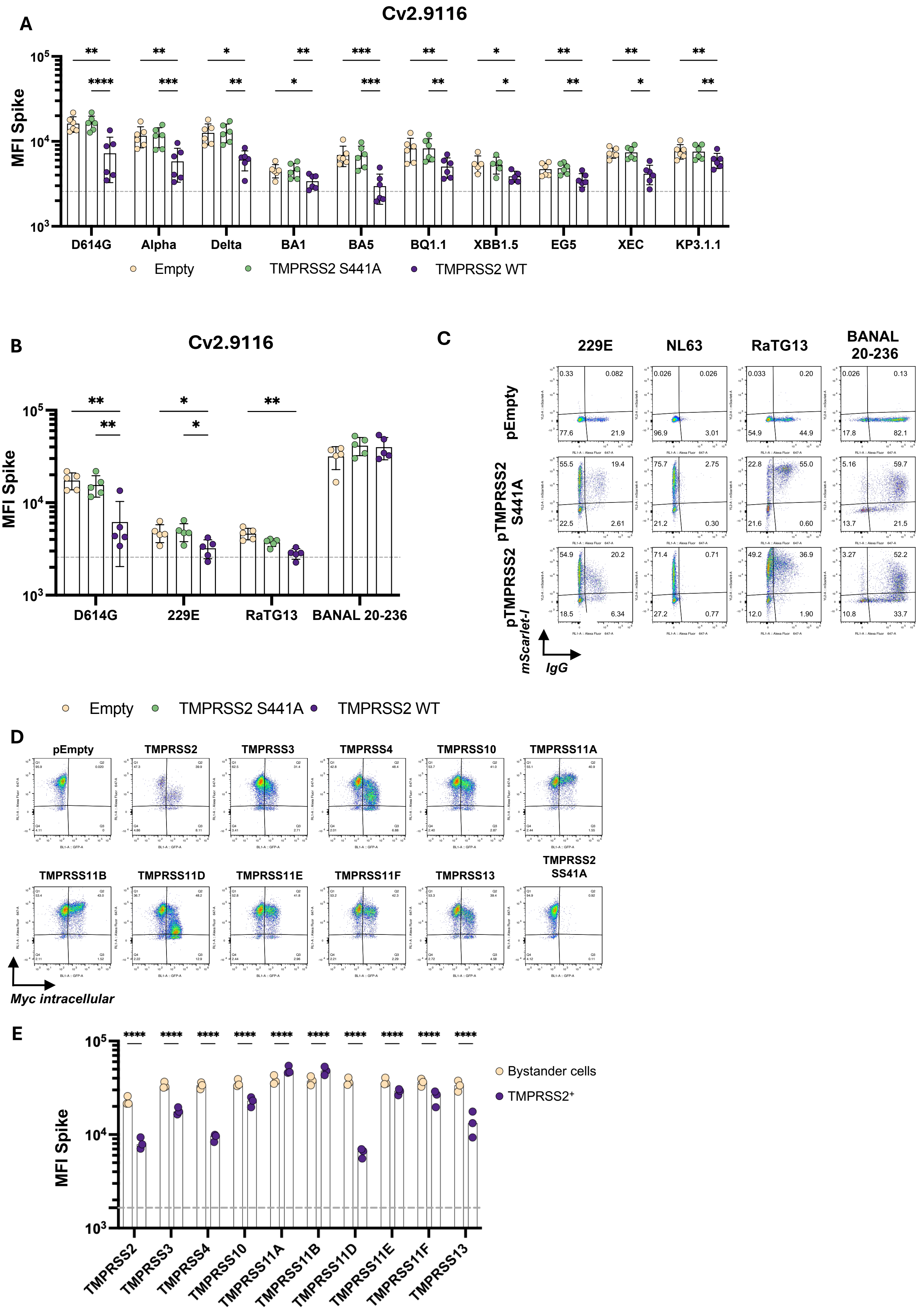

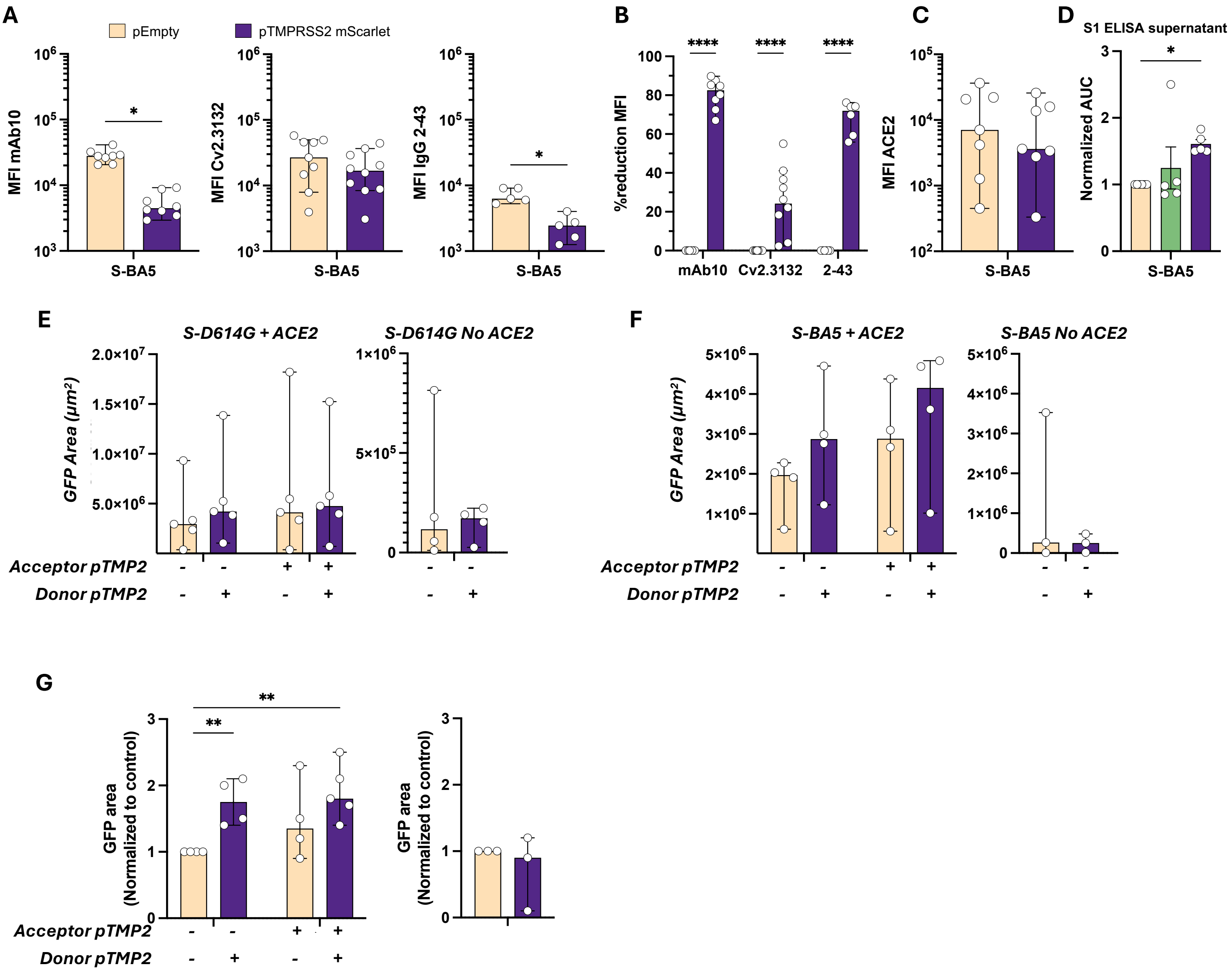

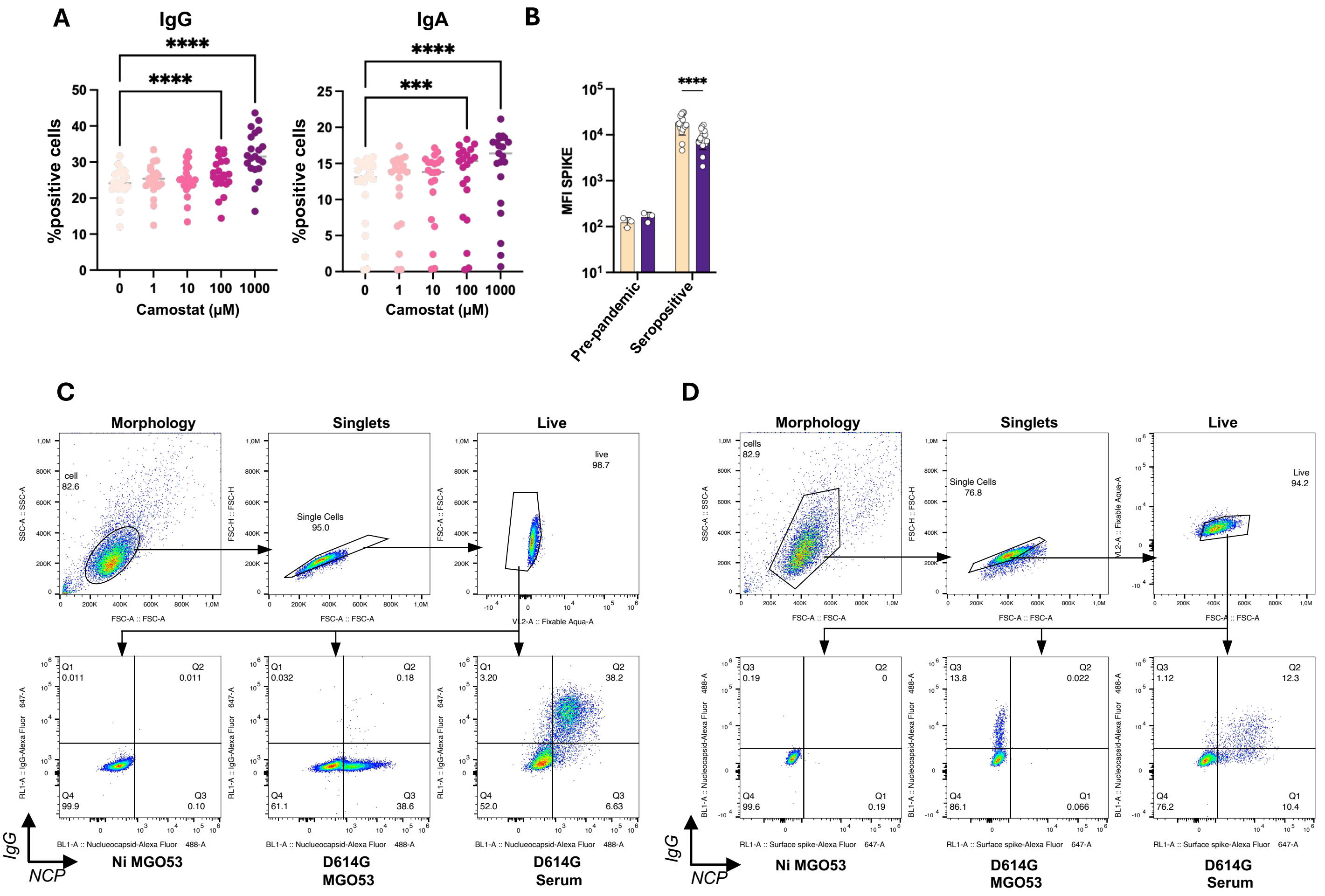

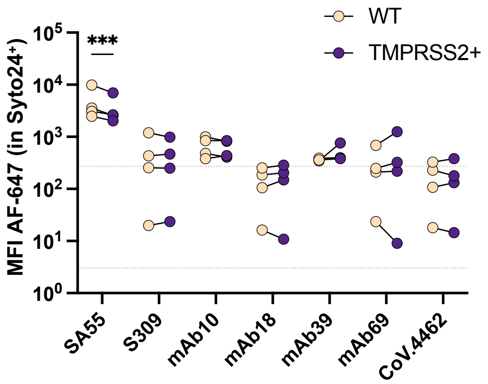
